# A host–gut microbial co-metabolite of aromatic amino acids, *p*-cresol glucuronide, promotes blood–brain barrier integrity *in vivo*

**DOI:** 10.1101/2022.01.11.475932

**Authors:** Andrew V. Stachulski, Tobias B-A Knausenberger, Sita N. Shah, Lesley Hoyles, Simon McArthur

**Author notes:** Correspondence to Dr Andrew V. Stachulski, Prof Lesley Hoyles or Dr Simon McArthur. Supplementary Materials are available from figshare: https://figshare.com/projects/p-Cresol_glucuronide_promotes_blood_brain_barrier_integrity_in_vivo/130088.

## Abstract

**Purpose:** The sequential activity of gut microbial and host processes can exert a powerful modulatory influence on dietary components, as exemplified by the metabolism of the amino acids tyrosine and phenylalanine to *p*-cresol by gut microbes, and then to *p*-cresol glucuronide (pCG) by host enzymes. Although such glucuronide conjugates are classically thought to be biologically inert, there is accumulating evidence that this may not always be the case. We investigated the activity of pCG, studying its interactions with the cerebral vasculature and the brain *in vitro* and *in vivo*.

**Methods:** Male C57Bl/6J mice were used to assess blood–brain barrier (BBB) permeability and whole brain transcriptomic changes in response to pCG treatment. Effects were then further explored using the human cerebromicrovascular endothelial cell line hCMEC/D3, assessing paracellular permeability, transendothelial electrical resistance and barrier protein expression.

**Results:** Mice exposed to pCG showed reduced BBB permeability and significant changes in whole brain transcriptome expression. Surprisingly, treatment of hCMEC/D3 cells with pCG had no notable effects until co-administered with bacterial lipopolysaccharide, at which point it was able to prevent the permeabilising effects of endotoxin. Further analysis suggested that pCG acts as an antagonist at the principal lipopolysaccharide receptor TLR4.

**Conclusion:** The amino acid phase II metabolic product pCG is biologically active at the BBB, highlighting the complexity of gut microbe to host communication and the gut–brain axis.

## Introduction

That communication between the gut microbiota and the brain can occur is now well established, with increasing evidence indicating a central role for microbe-derived metabolites acting primarily through three routes: directly on enteric and gut extrinsic neural pathways, by modification of enteroendocrine signalling or, as we and others have shown, via the circulation and interactions with the blood–brain barrier (BBB) [1, 2]. Notably, while many microbe-derived metabolites circulate in their native form, many others are subjected to host metabolic enzyme-mediated biotransformation, thereby altering their biological activities.

A good example of this lies in the metabolism of the microbial product *p*-cresol. This molecule is produced by bacterial fermentation of dietary tyrosine and phenylalanine in the colon [3], and passes through the gut epithelium into the portal vasculature. Notably, *p*-cresol undergoes extensive conjugation both in enterocytes [4] and by hepatic enzymes upon reaching the liver [5], such that it is found as, predominantly, *p*-cresol sulfate (pCS) and *p*-cresol glucuronide (pCG) in the systemic circulation [6, 7]. While pCS has been extensively studied in light of its role as a major uraemic toxin [6], the potential biological actions of pCG have received far less attention.

Classically, glucuronidation is considered as part of the phase II metabolic pathways, with the actions of the numerous UDP-glucuronosyltransferases serving to enhance renal clearance of parent compounds [8]. More recent evidence suggests that this form of conjugation may not always be a neutralising process however, with a number of clinically relevant molecules, including morphine, codeine and ethanol being known to gain pharmacological activity upon glucuronide conjugation [9–11]. Whether the same can be said for microbe-derived compounds present in the circulation is unclear, with studies into this question hindered by difficulties in obtaining pure molecules for study. We have recently established a novel pathway for chemical synthesis of pCG, and here we employ a combined *in vitro*/*in vivo* approach to identify the actions of this compound on the cerebral vasculature and the brain.

## Materials & Methods

### Drugs & Reagents

Trimethylsilyltrifluoromethanesulfonate was purchased from Fluorochem Ltd. UK and methyl 1,2,3,4-tetra-O-acetyl-β-D-glucuronate from Carbosynth UK. Solvents were of minimum HPLC grade and were purchased from Fisher Scientific UK. Ultrapure lipopolysaccharide (LPS) from *Porphyromonas gingivalis* was purchased from InvivoGen (Toulouse, France). Evans blue, 70 kDa FITC-dextran and MTT (3-(4,5-dimethylthiazol-2-yl)-2,5-diphenyltetrazolium bromide) were purchased from Merck Life Science UK Ltd., UK.

### Animals

Wild-type male C57Bl/6J mice aged between 7 and 8 weeks (Charles River UK Ltd., Margate, UK) were used for all experiments. Mice were kept under a 12 h:12 h light:dark regime, with *ad libitum* access to standard chow and drinking water; all animals were acclimatised to the holding facility environment for one week prior to experimentation. Animals were treated as described below and killed by transcardial perfusion with ice-cold saline under pentobarbitone anaesthesia. All experiments were approved by the QMUL Animal Welfare and Ethical Review Board and were performed in accordance with the UK Animals (Scientific Procedures) Act, 1986, under Project Licence PFA5C4F4F.

### In vivo BBB permeability analysis

Mice (n=5-6 per group) were injected i.p. with 1 mg/kg body weight pCG in 100 μl saline vehicle, a dose calculated to approximately double circulating concentrations [12], followed 2 h or 6 h later by assessment of Evans blue extravasation. One hour before assessment animals were injected i.p. with 100 μl of a 2% (w/v) solution of Evans blue dye in 0.9 % saline. Dye was permitted to circulate for 1 h before animals were transcardially perfused with 0.9% saline at 4 °C to remove dye remaining in the vasculature. Blood samples were allowed to coagulate at 37 °C for 15 minutes prior to centrifugation at 800 *g* for 10 minutes to separate serum. Brains were removed and homogenized by maceration in 0.1 M phosphate-buffered saline. Suspended macromolecules were precipitated by incubation with 60% trichloroacetic acid, and dye content of resulting supernatants was detected using a CLARIOstar spectrophotometer (BMG Labtech GmbH, Germany) alongside a standard curve of defined concentrations of Evans blue in the same buffer. Brain Evans’ blue content was expressed as μg of dye per mg of brain tissue, normalized to circulating serum concentrations.

### RNAseq data analyses

Processing of mouse brain samples (taken at 2 h) and RNA extraction were performed as described previously [2]. RNA samples (*n*=6 pCG, *n*=6 control) were sent to Macrogen Inc. (Seoul, Republic of Korea) where they were subject to quality checks (RIN analysis); libraries were prepared (TruSeq Stranded mRNA LT Sample Prep Kit) for paired-end (2x 100 nt) sequencing on an Illumina HiSeq 4000 apparatus. Three pCG-treated and three saline controls produced data of poor quality so were excluded from analyses after quality checks and consideration of Macrogen quality reports; as such, samples SG1, SG3, SG5, CG2, CG5 and CG6 were used for all analyses described from hereon. Raw RNAseq (fastq) sequence data were processed in house as described previously [2]. Entrez gene identifiers were converted to gene symbols using *Mus musculus* annotations downloaded from NCBI on 4 January 2021; only those genes with valid Entrez gene identifiers were retained in analyses. Significantly differentially expressed genes (FDR *P*<0.05) identified using DESeq2 v1.22.1 [13] were analysed by mouse KEGG pathway over-representation analysis using Enrichr [14, 15] and manual curation. Signaling Pathway Impact Analysis (SPIA v v1.22.1) [16] was used to determine whether Kyoto Encyclopedia of Genes and Genomes (KEGG) *Mus musculus* pathways (downloaded on 22 December 2021) were activated or inhibited in mouse brain cells exposed to pCG. RNAseq data have been deposited in ArrayExpress under accession number E-MTAB-11340. Normalized and log2-transformed RNAseq data are available as Supplementary Material (Supplementary Table 1).

### Cell culture

The human cerebromicrovascular endothelial cell line hCMEC/D3 was maintained and treated as described previously [17]. Cells were cultured to confluency in complete endothelial cell growth medium MV2 (PromoCell GmbH, Germany), whereupon VEGF was removed and cells were further cultured for a minimum of 4 days to enable intercellular tight junction formation prior to experimentation. All cell cultures were used below passage 35 to ensure retention of appropriate endothelial characteristics [18].

### Cell survival analysis

The potential for pCG-induced cytotoxicity was assessed using the MTT assay. Briefly, cells were treated with pCG for 24 h, prior to administration of MTT at 500 μg/ml. Cells were incubated at 37 °C for 2 h, medium was removed and resulting crystals were solubilised by incubation for 2 minutes in dimethyl sulfoxide. Absorbance was read at 540 nm using a CLARIOstar spectrophotometer (BMG Labtech, Ortenberg, Germany), with a reference wavelength at 570 nm.

### In vitro barrier function assessments

Paracellular permeability and transendothelial electrical resistance were measured on 100% confluent cultures polarised by growth on 24-well plate polyethylene terephthalate (PET) transwell inserts (surface area: 0.33 cm^2^, pore size: 0.4 μm; Appleton Woods, UK) previously coated with calf-skin collagen (15 μg/cm^2^ and fibronectin 3 μg/cm^2^; both Merck Life Science UK Ltd.). The permeability of hCMEC/D3 cell monolayers to 70 kDa FITC-dextran (2 mg/ml) was measured as described previously [19, 20]. Transendothelial electrical resistance (TEER) measurements were performed using a Millicell ERS-2 Voltohmmeter (Millipore, Watford, UK) and were expressed as Ω.cm^2^. In all cases, values obtained from cell-free inserts similarly coated with collagen and fibronectin were subtracted from the total values.

### Immunofluorescence

Confluent hCMEC/D3 monolayers grown on transwell inserts as described above were fixed by immersion in 2% formaldehyde in 0.1 M PBS for 10 minutes at room temperature. Cells were immunostained according to standard protocols [21] using a primary rabbit anti-human antibody directed against zona occludens-1 (ZO-1; 1:100, Thermo-Fisher Scientific, UK) and a AF488-conjugated secondary goat anti-rabbit antibody (1:500, ThermoFisher Scientific, UK) or AF488-conjugated phalloidin (100 nM; Cytoskeleton Inc., Denver, USA). Nuclei were counterstained with DAPI (50 ng/ml; Merck Life Science UK Ltd., UK). Images were captured using an LSM880 confocal laser scanning microscope (Carl Zeiss Ltd., Cambridge, UK) fitted with 405 nm and 488 nm lasers and a 63x oil immersion objective lens (NA, 1.4 mm, working distance, 0.17 mm). Images were captured with ZEN imaging software (Carl Zeiss Ltd., UK) and analysed using ImageJ 1.53c (National Institutes of Health, USA).

### Flow cytometry

Cells were labelled with APC-conjugated mouse monoclonal anti-CD11b (Biolegend, UK), FITC-conjugated mouse monoclonal anti-CD14 (Biolegend, UK), FITC-conjugated mouse monoclonal anti-MD2 (Biolegend, UK), PE-conjugated mouse monoclonal anti-TLR4, APC-conjugated mouse monoclonal anti-BCRP (BD Biosciences, Oxford, UK), or PE-conjugated mouse monoclonal anti-MDR1A (BD Biosciences, UK), for analysis by flow cytometry. Briefly, cells were treated as described below and, in the case of hCMEC/D3 cells, detached using 0.05% trypsin and incubated with antibodies for 20 minutes at 4 °C. Immunofluorescence was analysed for 10,000 events per treatment using a BD FACS Canto II flow cytometer (BD Biosciences,UK), and data were analysed using FlowJo 8.0 software (Treestar Inc., CA, USA).

### Efflux transporter assays

Activity of the major efflux transporters P-glycoprotein and BCRP was determined using commercially available assays (PREDEASY™ ATPase Assay Kits, Solvo Biotechnology Inc., Budapest, Hungary), performed according to the manufacturer’s instructions. Stepwise dose– response curves centred around reported physiological circulating concentrations of pCG (12.3 nM – 27 μM) were constructed (n = 4) to investigate stimulatory and inhibitory effects upon transporter activity.

### Statistical analysis

Sample sizes were calculated to detect differences of 15% or more with a power of 0.85 and α set at 5%, calculations being informed by previously published data [2, 17]. Experimental data are expressed as mean ± SEM, with n = 6-9 independent experiments for all studies. In all cases, normality of distribution was established using the Shapiro–Wilk test, followed by analysis with two-tailed Student’s *t*-tests to compare two groups or, for multiple comparison analysis, one- or two-way ANOVA followed by *post hoc* analysis by either Dunnett’s test (for dose-response experiments) or Tukey’s HSD test (all other comparisons). A P value of less than or equal to 5% was considered significant.

## Results

### Synthesis of p-cresol glucuronide

As previously described [22] and shown schematically (Fig. 1A), reaction of *p*-cresol **1** with 1,2,3,4-tetra-O-acetyl-β-D-glucuronate **2** in CH2Cl2 promoted by trimethylsilyltrifluoromethanesulfonate afforded the conjugate **3** in very good yield as a single β-anomer. Hydrolysis of **3** under mild conditions (aq. Na2CO3, MeOH) afforded the desired glucuronide sodium salt **4** after partial neutralisation to pH 6. Recrystallisation gave material of microanalytical purity, as indicated by the ^1^H NMR spectrum (Fig. 1B).

**Figure 1:**
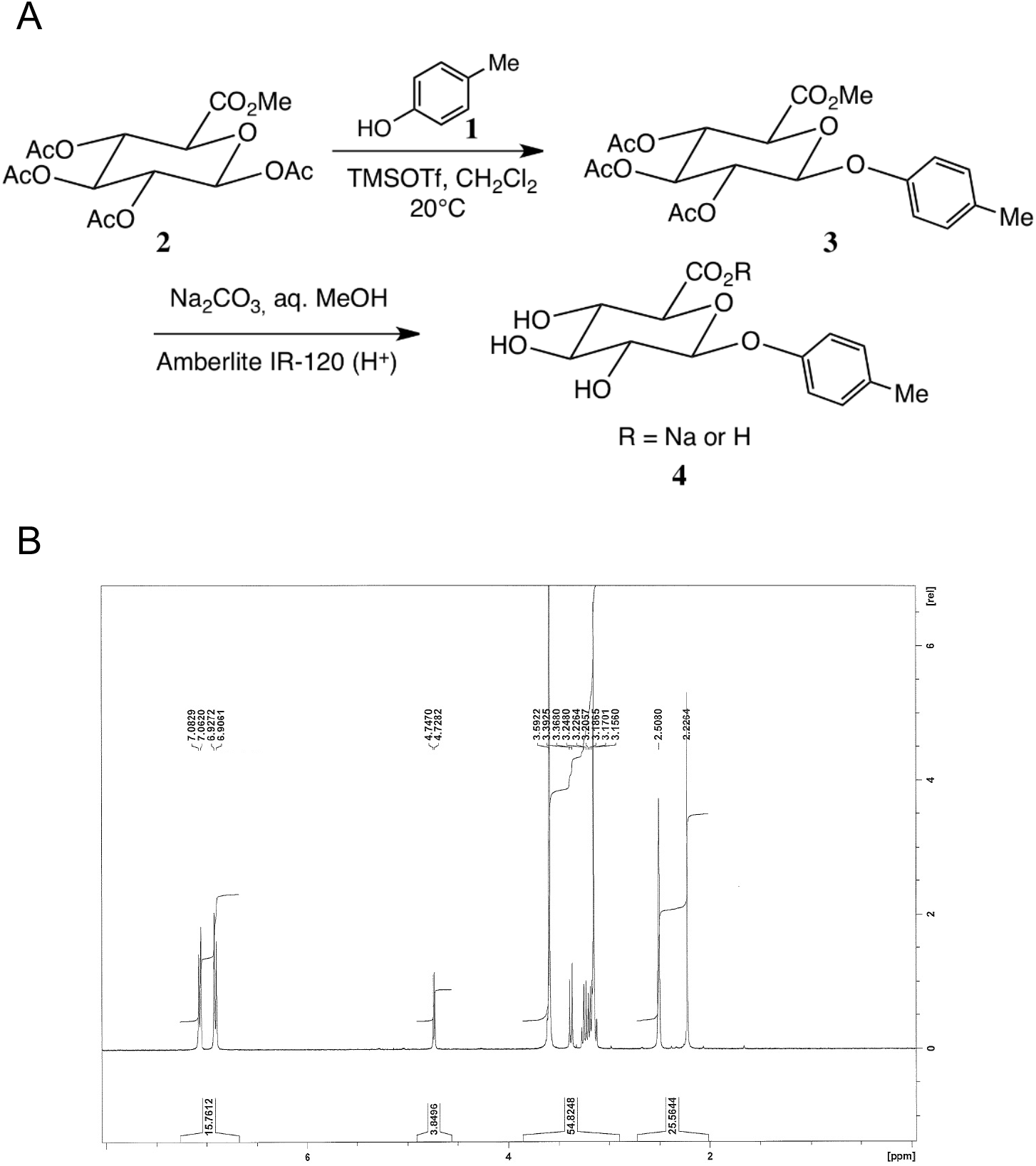
Production and validation of pCG. **A)** Schematic synthetic pathway for the production of pCG, as previously reported [22]. **B)** Typical [^1^H]-NMR spectroscopy trace indicating microanalytical purity of *de novo* synthesised pCG.

### pCG modulates BBB integrity and the whole brain transcriptome in vivo

A defining property of the cerebral vasculature is the existence of a tight barrier function limiting passage of soluble molecules into the brain parenchyma, the BBB. We examined whether exposure to increased levels of pCG could affect BBB integrity *in vivo*, assessed by monitoring extravasation of administered Evans blue dye into the brain tissue. Treatment of mice with 1 mg/kg pCG i.p. (a dose known to approximately double baseline serum concentrations [23]) caused a significant reduction in entry of Evans blue to the brain tissue, by approximately 50% within 6 h of treatment (Fig. 2A).

**Figure 2:**
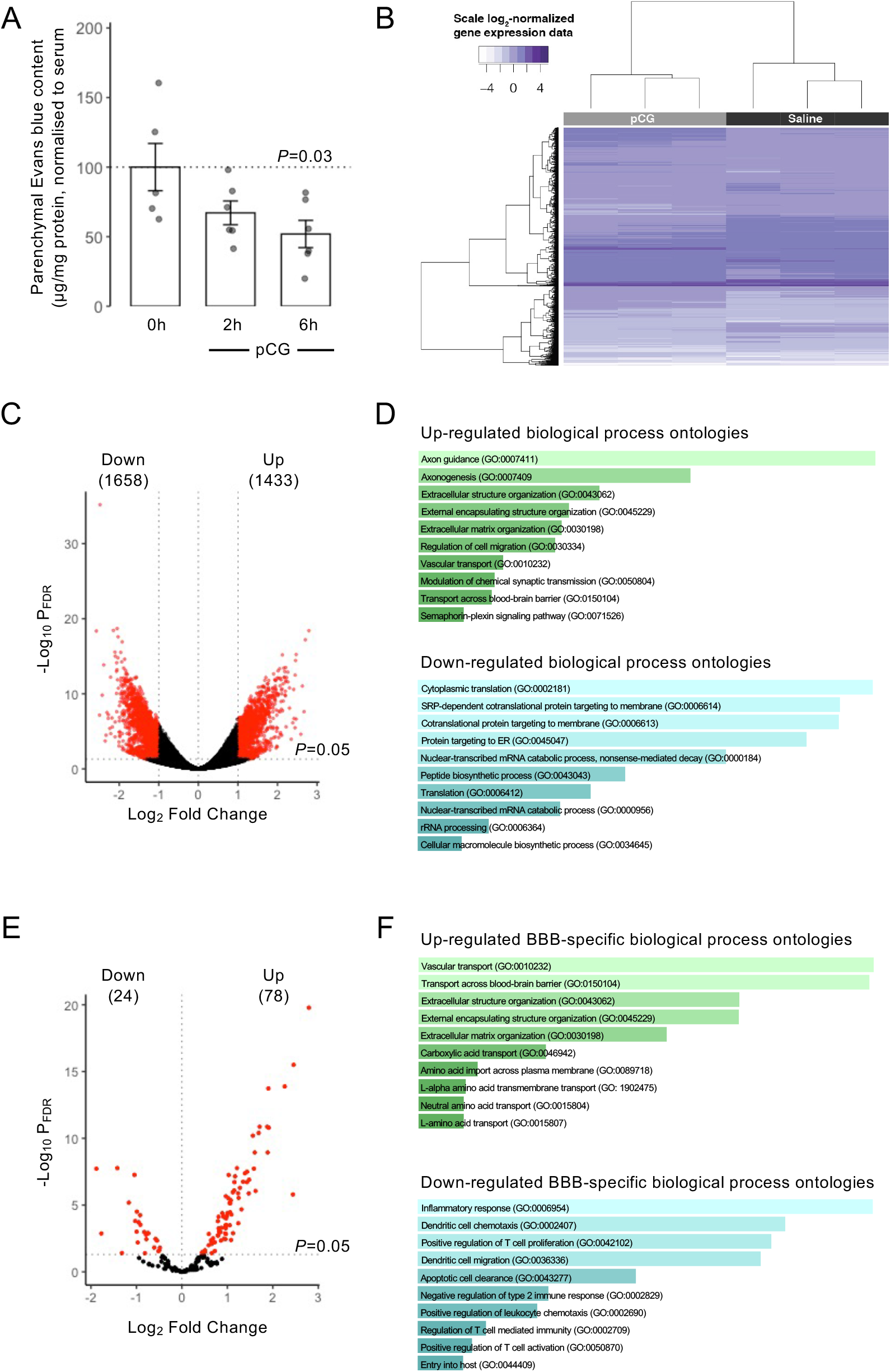
pCG treatment alters murine BBB permeability and CNS transcriptional profile *in vivo*. **A)** Treatment of male C57Bl/6 mice by i.p. injection of pCG (1 mg/kg) caused a time-dependent reduction in extravasation of Evans blue tracer into the CNS parenchyma, reaching statistical significance 6 h post administration; data are mean ± s.e.m. n=6. **B)** Heatmap showing expression of the 7702 genes found to be significantly (PFDR<0.05) differentially expressed in the CNS of male C57Bl/6 mice 2 h following i.p. injection of 1 mg/kg pCG (n=3 per group). **C)** Volcano plot showing 3091 significantly (PFDR< 0.05) 2-fold differentially expressed genes (red dots). **D)** Biological processes associated with genes found to be significantly and ≥ 2-fold upregulated (n=1433) or downregulated (n=1658) upon exposure of mice to pCG. Images are based on Enrichr P value ranking from GO analysis, the lighter the colour and longer the bar, the more is significant is the result, as determined by rank-based ranking; only the top 10 results are shown in each case. **E)** Volcano plot showing significantly (PFDR< 0.05) differentially expressed BBB-relevant genes (red dots). **F)** Biological processes associated with BBB-relevant genes found to be significantly upregulated (n=78) or downregulated (n=24) upon exposure of mice to pCG. Images are based on Enrichr P value ranking from GO analysis, the lighter the colour and longer the bar, the more is significant is the result, as determined by rank-based ranking; only the top 10 results are shown in each case.

To investigate the mechanism(s) underlying this action of pCG, we performed bulk RNAseq analysis of brain tissue from animals treated for 2 h with 1 mg/kg pCG i.p., identifying 7702 significantly differentially expressed genes (Fig. 2B; Supplementary Table 2), of which 1658 and 1433 showed greater than 2-fold up- or down-regulation respectively following correction for multiple testing (Fig. 2C). Analysis of gene ontology categories over-represented within these gene sets using Enrichr [14, 15, 24] identified a number of different biological process ontologies exhibiting significant changes (Fig. 2D), with ontologies relating to axon generation and extracellular matrix organisation being notably up-regulated, while pathways associated with protein synthesis and ribosomal activity were down-regulated. SPIA of all differentially expressed genes revealed several significantly over-represented KEGG pathways (Suppl. Fig. 2A), notably indicating pathways associated with growth factor/transcription factor signalling and the response to infection as being activated, whilst pathways associated with cellular degradation and metabolism were inhibited (Suppl. Fig. 2B).

To specifically examine the interactions of pCG with the BBB, we further interrogated transcriptomic changes induced by pCG treatment by comparison with a defined list of 203 known BBB-relevant genes [17], identifying a total of 78 upregulated and 24 down-regulated number of genes exhibiting statistically significant regulation (Fig. 2E; Supplementary Table 3). Examination of associated biological process gene ontologies here identified clear upregulation in multiple transport pathways and suppression of inflammatory processes (Fig. 2F). Individual gene-level analysis of differentially expressed transporter systems identified enhanced expression of a wide range of nutrient uptake transporters, whereas in contrast only the transporters for myo-inositol and transferrin and aquaporin-4 were significantly down-regulated (Table 1).

**Table 1:** Significantly differentially expressed BBB-associated transporter genes following pCG treatment *in vivo*

| Gene | Name | Saline | pCG | Direction | P<sub>FDR</sub> |
| --- | --- | --- | --- | --- | --- |
| *Ldlr* | Low density lipoprotein receptor | 6.91 ± 0.32 | 8.67 ± 0.42 | Up | 3.98 x 10<sup>-11</sup> |
| *Abca2* | ATP binding cassette subfamily A member 2 | 10.49 ± 0.77 | 12.61 ± 0.42 | Up | 1.15 x 10<sup>-9</sup> |
| *Slc2a1* | Solute carrier family 2 member 1 | 9.95 ± 0.48 | 11.66 ± 0.35 | Up | 1.15 x 10<sup>-9</sup> |
| *Slc38a3* | Solute carrier family 38 member 3 | 9.36 ± 0.47 | 10.84 ± 0.32 | Up | 4.15 x 10<sup>-8</sup> |
| *Slc1a4* | Solute carrier family 1 member 4 | 9.28 ± 0.21 | 10.48 ± 0.28 | Up | 6.98 x 10<sup>-8</sup> |
| *Slc7a5* | Solute carrier family 7 member 5 | 8.94 ± 0.6 | 10.55 ± 0.37 | Up | 1.26 x 10<sup>-7</sup> |
| *Slc6a9* | Solute carrier family 6 member 9 | 8.40 ± 0.63 | 9.98 ± 0.32 | Up | 1.92 x 10<sup>-7</sup> |
| *Slc7a1* | Solute carrier family 7 member 1 | 8.54 ± 0.34 | 9.75 ± 0.28 | Up | 8.06 x 10<sup>-7</sup> |
| *Slc38a5* | Solute carrier family 38 member 5 | 4.61 ± 0.5 | 6.42 ± 0.23 | Up | 8.78 x 10<sup>-7</sup> |
| *Slc27a4* | Solute carrier family 27 member 4 | 9.73 ± 0.27 | 10.88 ± 0.31 | Up | 1.10 x 10<sup>-6</sup> |
| *Slc16a2* | Solute carrier family 16 member 2 | 8.35 ± 0.34 | 9.62 ± 0.44 | Up | 1.30 x 10<sup>-6</sup> |
| *Slc22a8* | Solute carrier family 22 member 8 | 7.65 ± 0.37 | 8.77 ± 0.21 | Up | 7.13 x 10<sup>-6</sup> |
| *Slc29a4* | Solute carrier family 29 member 4 | 7.01 ± 0.73 | 8.50 ± 0.45 | Up | 2.09 x 10<sup>-5</sup> |
| *Mfsd2a* | Major facilitator superfamily domain containing 2A | 8.51 ± 0.48 | 9.57 ± 0.21 | Up | 9.88 x 10<sup>-5</sup> |
| *Slc38a1* | Solute carrier family 38 member 1 | 10.63 ± 0.27 | 11.46 ± 0.11 | Up | 1.60 x 10<sup>-4</sup> |
| *Abcc4* | ATP binding cassette subfamily C member 4 | 7.20 ± 0.31 | 8.20 ± 0.49 | Up | 3.25 x 10<sup>-4</sup> |
| *Slco2b1* | Solute carrier organic anion transporter family member 2B1 | 7.93 ± 0.62 | 9.02 ± 0.22 | Up | 3.28 x 10<sup>-4</sup> |
| *Slc27a1* | Solute carrier family 27 member 1 | 9.77 ± 0.53 | 10.74 ± 0.15 | Up | 5.02 x 10<sup>-4</sup> |
| *Abcc1* | ATP binding cassette subfamily C member 1 | 8.00 ± 0.44 | 8.90 ± 0.25 | Up | 8.66 x 10<sup>-4</sup> |
| *Slc5a6* | Solute carrier family 5 member 6 | 8.21 ± 0.33 | 8.94 ± 0.23 | Up | 3.12 x 10<sup>-3</sup> |
| *Slc1a5* | Solute carrier family 1 member 5 | $5.11 \pm 0.36$ | $6.15 \pm 0.11$ | Up | $3.27 \times 10^{-3}$ |
| *Slc29a1* | Solute carrier family 29 member 1 (Augustine blood group) | $7.76 \pm 0.35$ | $8.45 \pm 0.08$ | Up | $4.39 \times 10^{-3}$ |
| *Slc6a6* | Solute carrier family 6 member 6 | $10.48 \pm 0.58$ | $11.31 \pm 0.15$ | Up | $6.05 \times 10^{-3}$ |
| *Slc1a1* | Solute carrier family 1 member 1 | $10.59 \pm 0.31$ | $11.13 \pm 0.13$ | Up | $2.41 \times 10^{-2}$ |
| *Slc44a1* | Solute carrier family 44 member 1 | $9.99 \pm 0.2$ | $10.44 \pm 0.23$ | Up | $4.07 \times 10^{-2}$ |
| *Slc5a3* | Solute carrier family 5 member 3 | $9.50 \pm 0.4$ | $8.55 \pm 0.37$ | Down | $2.99 \times 10^{-4}$ |
| *Tfrc* | Transferrin receptor | $11.39 \pm 0.29$ | $10.63 \pm 0.25$ | Down | $1.29 \times 10^{-3}$ |
| *Aqp4* | Aquaporin 4 | $11.83 \pm 0.32$ | $10.98 \pm 0.51$ | Down | $3.58 \times 10^{-3}$ |

### pCG has limited direct effects upon an in vitro model of the BBB

Following these *in silico* analyses, we sought to investigate the biological pathway(s) through which pCG affected the BBB, using a well-established model of the human brain capillary endothelium, the hCMEC/D3 cell line [25]. Initial assessment of potential pCG toxicity using the MTT assay showed no effects on cell survival following 24 h treatment of hCMEC/D3 cells with concentrations of up to 100 μM pCG (Fig. 3A). Similarly, as β-glucuronidase is known to be present in the cerebral endothelium, albeit at low levels [26], it is plausible that the effects of pCG may be caused by reversion to its parent *p*-cresol molecule. However, exposure of hCMEC/D3 cells to *p*-cresol itself (5 μM, 24 h) caused a significant increase in paracellular permeability to a 70 kDa FITC-dextran tracer and accompanying reduction in transendothelial electrical resistance (Suppl. Fig. 1), indicating an abrogation of BBB integrity, essentially the opposite of our *in vivo* findings.

**Figure 3:**
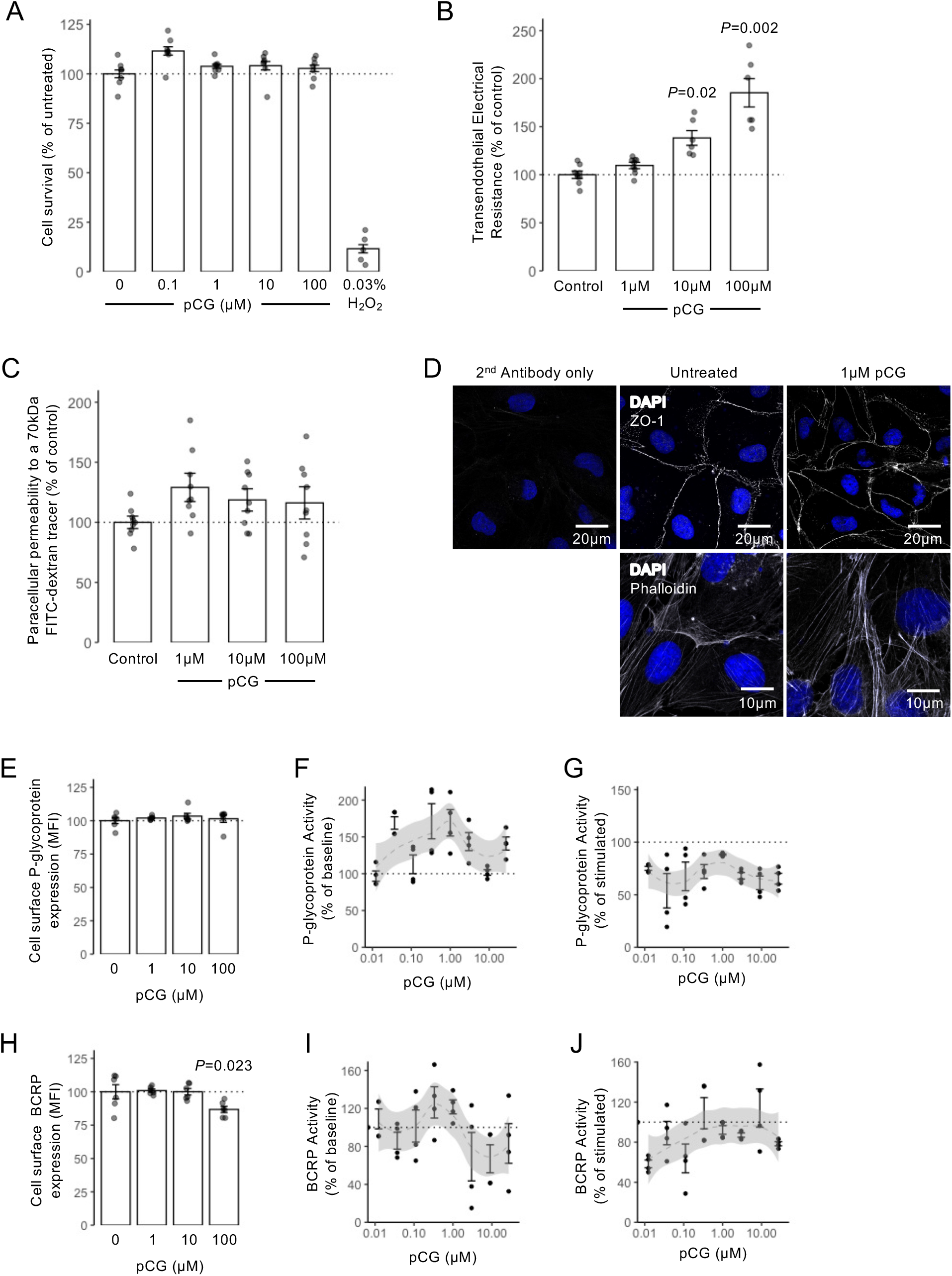
Limited effects of pCG upon unstimulated *in vitro* models of the BBB. **A)** Treatment of hCMEC/D3 cells with increasing doses of pCG (0.1 – 100 μM; 24 h) has no effect on cell survival or proliferation as measured by the MTT assay, in contrast to the highly toxic effects of 0.03% H2O2 exposure; data are mean ± s.e.m., n=4. **B)** Trans-endothelial electrical resistance across polarised hCMEC/D3 monolayers following 24 h treatment with pCG; data are mean ± s.e.m., n=6. **C)** Paracellular permeability of polarised hCMEC/D3 monolayers to a 70 kDa FITC-dextran tracer following 24 h treatment with pCG; data are mean ± s.e.m., n=9. **D)** Confocal microscopic analysis of expression of the tight junction component zona occludens-1 (ZO-1) or AF488-phalloidin labelled actin filaments in hCMEC/D3 cells following treatment for 24 h with 1 μM pCG. Images are representative of at least three independent experiments. **E)** Treatment of hCMEC/D3 cells with pCG (24 h) had no effect on cell surface expression of P-glycoprotein, data are mean ± s.e.m. n=6. **F, G)** Lack of stimulatory (F) or inhibitory (G) effects of pCG upon baseline or stimulated P-glycoprotein activity, data are mean ± s.e.m., n=4. **H)** Treatment of hCMEC/D3 cells with pCG (24 h) caused a slight but significant reduction in BCRP expression at the highest dose tested (100 μM), data are mean ± s.e.m., n=6. **I-J)** Lack of stimulatory (I) or inhibitory (J) effects of pCG upon baseline or stimulated P-glycoprotein activity, data are mean ± s.e.m., n=4.

We then examined the ability of pCG itself to affect hCMEC/D3 monolayer barrier integrity. Exposure of hCMEC/D3 cells for 24 h to pCG caused a dose-dependent increase in transendothelial electrical resistance, becoming statistically significant with 10 μM and 100 μM concentrations (Fig. 3B), but this was not accompanied by any change in permeability to the 70 kDa FITC-dextran tracer (Fig. 3C). Microscopic examination of the tight junction component ZO-1 and the actin cytoskeleton similarly revealed little effect of pCG upon the endothelial cells (Fig. 3D).

As our transcriptomic data indicated upregulation of multiple nutrient uptake transporter genes, we investigated whether pCG could also affect two of the principal efflux transport systems of the BBB, namely P-glycoprotein and BCRP. While pCG had no effect on cell surface P-glycoprotein expression at any dose tested (Fig. 3E), exposure of cells to 100 μM pCG did cause a slight, but significant reduction in BCRP expression (Fig. 3H). Neither transporter was activated or inhibited by the presence of pCG at any concentrations tested (Fig. 3F-G, I-J).

### pCG antagonises the BBB-permeabilising actions of bacterial LPS

In light of the contrast between the limited effects of pCG seen in our *in vitro* BBB model, we sought alternative explanations for the more pronounced effects of the metabolite seen *in vivo*, taking a lead from the indicated suppression of inflammatory process ontologies. Several structurally dissimilar glucuronidated molecules interact with the bacterial LPS receptor TLR4 and its heterodimeric partner MD-2, including morphine-3-glucuronide [27], ethyl glucuronide [9] and a range of steroid hormone glucuronide conjugates [28], leading us to hypothesise that this may also be the case for pCG. LPS is known to circulate at low, but non-zero, levels in normal mice and humans [29, 30], and is known to enhance BBB permeability *in vitro* and *in vivo* [2], hence we investigated the interaction between it and pCG in our model system.

We initially confirmed that hCMEC/D3 cells express TLR4 and its accessory proteins MD-2 and CD14 (Suppl. Fig. 3A-C). Treatment of hCMEC/D3 cells with LPS (*Porphyromonas gingivalis*, 10 ng/ml, 24 h) significantly enhanced paracellular permeability to a 70 kDa FITC-dextran tracer (Fig. 4A) and reduced transendothelial electrical resistance (Fig. 4B), effects that were both prevented by 30 minutes pre-treatment with pCG (1 μM). Similar treatment of endothelial monolayers with LPS disrupted circumferential localisation of the key tight junction molecule ZO-1 (Fig. 4C) and induced the appearance of large numbers of cytosolic actin fibres (Fig. 4D), both of which features were prevented by 30 minutes pre-treatment with pCG (1 μM). This effect did not appear to be due to down-regulation of TLR4 or its accessory molecules CD14 or MD-2 on the surface of the endothelial cells (Fig. 4E-G), suggesting pCG may be acting as an antagonist at this receptor.

**Figure 4:**
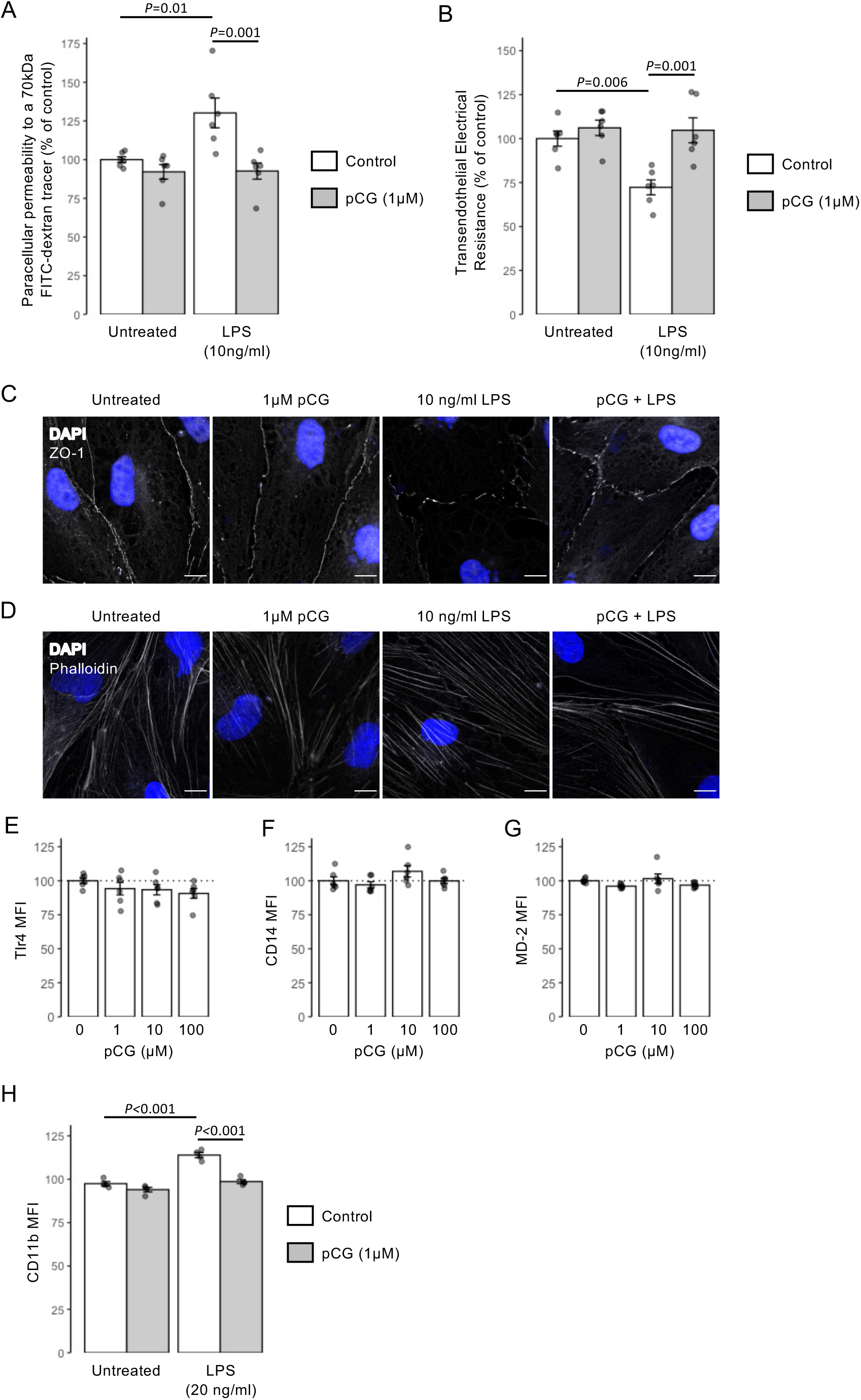
Treatment with pCG antagonises the effects of LPS *in vitro*. **A)** Paracellular permeability of polarised hCMEC/D3 monolayers to a 70 kDa FITC-dextran tracer with or without 24 h treatment with *Porphyromonas gingivalis* LPS (10 ng/ml) under control conditions or with 30 minutes pre-treatment with 1 μM pCG; data are mean ± s.e.m., n=6. **B)** Transendothelial electrical resistance of polarised hCMEC/D3 monolayers to a 70 kDa FITC-dextran tracer with or without 24 h treatment with *P. gingivalis* LPS (10 ng/ml) under control conditions or with 30 minutes pre-treatment with 1 μM pCG; data are mean ± s.e.m., n=6. **C)** Confocal microscopic analysis of expression of the tight junction component zona occludens-1 (ZO-1) in hCMEC/D3 cells following treatment for 24 h with 10 ng/ml LPS with or without 30 minutes prior administration of 1 μM pCG. Images are representative of at least three independent experiments. **D)** Confocal microscopic analysis of AF488-phalloidin defined actin filaments in hCMEC/D3 cells following treatment for 24 h with 10 ng/ml LPS with or without 30 minutes prior administration of 1 μM pCG. Images are representative of at least three independent experiments. **E)** Treatment of hCMEC/D3 cells with pCG (1 – 100 μM, 24 h) has no effect on surface expression of TLR4; data are mean ± s.e.m., n=6. **F)** Treatment of hCMEC/D3 cells with pCG (1 – 100 μM, 24 h) has no effect on surface expression of CD14; data are mean ± s.e.m., n=6. **G)** Treatment of hCMEC/D3 cells with pCG (1 – 100 μM, 24 h) has no effect on surface expression of MD-2; data are mean ± s.e.m., n=6. **H)** Pre-treatment for 30 minutes with pCG (1 μM) prevents the increase in cell surface CD11b expression on THP-1 monocyte-like cells induced by 24 h treatment with LPS (20 ng/ml); data are mean ± s.e.m., n=4.

To provide further support for this hypothesis, we investigated whether pCG could functionally antagonise an alternative and unrelated effect of LPS treatment, upregulation of surface expression of the integrin CD11b on the human monocyte cell line THP-1 (Suppl. Fig. 3). Treatment of THP-1 cells with LPS (20 ng/ml, 24h) significantly up-regulated surface CD11b expression, an effect prevented by 30 minutes pre-treatment with 1 μM pCG (Fig. 4G), confirming the ability of pCG to antagonise LPS-induced signalling responses in distinct circumstances.

## Discussion

Glucuronidation is a key stage in phase II metabolism and clearance of endogenous and exogenous molecules and has long been investigated in this regard. Much is now known about the various UDP-glucuronosyltransferases responsible for glucuronidation at different sites in the body [8], but the biological actions of glucuronide compounds once they have been formed are rather less understood. In most cases, glucuronide conjugates have been considered as biologically inactive and simply destined for renal elimination, but our data add to the steadily building picture that this may not be universally true. Notably, glucuronide derivatives of morphine, ethanol and estradiol have been shown to act as agonists of the TLR4 complex, promoting allodynia and inflammation upon spinal cord administration [9, 27, 28]. Our data add the tyrosine/phenylalanine metabolite pCG to the list of glucuronide conjugates that can interact with TLR4 signalling, but with the marked difference that, in contrast to the other known activating agents, pCG is a functional antagonist and prevents the permeabilising effects of bacterial endotoxin exposure upon the BBB.

Whilst pCG has long been known to circulate in the blood, its physiological and potentially pathological actions have remained somewhat elusive. Our description of an antagonistic action of pCG upon the principal LPS receptor, the TLR4 complex, indicates an anti-inflammatory effect of the molecule and suggests that it may, at least at physiological concentrations, aid cerebrovascular resilience to the damaging effects of LPS exposure [21, 31], thereby protecting against the development of sickness behaviours [32]. However, pCG is also well known as a potential uraemic toxin [33]. Exposure at levels seen in patients undergoing haemodialysis has been reported to directly evoke a low level of endothelial reactive oxygen species release [34], to impair endothelial succinate dehydrogenase function [35] and to potentiate some of the inflammatory effects of pCS upon leukocytes [36] and the endothelium [37]. Notably, individuals with renal dysfunction have increased susceptibility to bacterial infection [38–40] despite the presence of chronic low grade leukocyte activation [41]. Given that the majority of circulating pCG in such patients is freely available [42] and thus presumably able to interact with TLR4, the potential contribution that such antagonism by pCG makes to masking signs of bacterial infection bears further investigation.

Beyond emphasising the need to look again at glucuronide conjugates as potential biological actors, our data also highlight the position of the cerebral vasculature and the BBB as targets for the actions of microbial metabolites and an important aspect of the gut–brain axis. A range of gut microbe-derived metabolites, including short-chain fatty acids, methylamines and, here, cresols, have now been shown to regulate BBB integrity [1, 2, 17] *in vivo*. That such structurally diverse molecules can modulate BBB function epitomises the complexity of the gut microbiome–brain axis, but also emphasises the importance of systematic investigation of this communication pathway. Moreover, as pCG is a product of both gut microbial and host enzymatic co-metabolism of aromatic amino acids, our data emphasise the need to consider both microbial and host systems in regulating gut microbe–brain communication. With over 200 known microbe-derived metabolites present in the human circulation [43], there is clearly much still to learn about how they might influence the cerebral vasculature and their implications for health, ageing and disease.

A notable feature of the gut microbiota is its exquisite sensitivity to dietary change [44], with the make-up of the gut microbiome changing in a matter of weeks of exposure to a novel diet [45]. As diet is also known to be a major risk factor for cerebrovascular and neurological health [46], studying the links between diet, gut microbe–host co-metabolites and the BBB may be instructive in understanding the pathogenesis of and, potentially, treatment for neurovascular disease. In particular, our study of the simple phenolic glucuronide pCG may be of relevance when it comes to understanding the actions of its more chemically complex relatives, the dietary polyphenol glucuronides. Diets supplemented with foods containing polyphenols have been shown to improve cerebral blood flow and neurovascular coupling in humans [47–51], and rodent studies have revealed polyphenols to protect against ischaemia or trauma-induced BBB integrity damage [52–55]. Notably, however, such dietary polyphenols are primarily found in the circulation as conjugates: sulfates, methylates, and, conspicuously, glucuronides [56]. At the least, the presence of high levels of glucuronide conjugates suggest that these agents should be investigated as potential mediators of the beneficial effects of dietary polyphenols upon the cerebral vasculature.

### Conclusion

Here, we show that pCG, thought to be a relatively inert product of gut microbe–host enzyme co-metabolism, can influence the BBB and potentially immune cell activity through functional antagonism at the TLR4 complex. This adds to our understanding of the role of glucuronide conjugates as not only targets for renal elimination, but also as potent biological actors in their own right. Moreover, our data emphasise the importance of considering both microbial and host metabolic processes in understanding the mechanism(s) of communication that underlie the gut microbiota–brain axis.

## Statements & Declarations

The authors have no competing interests to declare that are relevant to the content of this article. Author Contributions: AVS prepared and purified pCG, TBAK, SNS & SM performed cell culture and animal experiments, LH performed bioinformatic analyses. AVS, LH & SM wrote the manuscript. All authors have read and approved the final version of the manuscript.

## Acknowledgements

This work was funded by Alzheimer’s Research UK Pilot Grant No. ARUK-PPG2016B-6. PREDEASY™ efflux transporter analysis kits were generously provided through the SOLVO Biotechnology Research and Academic Collaborative Transporter Studies (ReACTS) Program. This work used the computing resources of the UK MEDical BIOinformatics partnership—aggregation, integration, visualisation and analysis of large, complex data (UK MED-BIO), which was supported by the Medical Research Council (grant number MR/L01632X/1). This project has received funding from the European Union’s Horizon 2020 research and innovation programme under grant agreement No 874583. This publication reflects only the authors’ view and the European Commission is not responsible for any use that may be made of the information it contains.

---

**Supplemental Figure 1:**
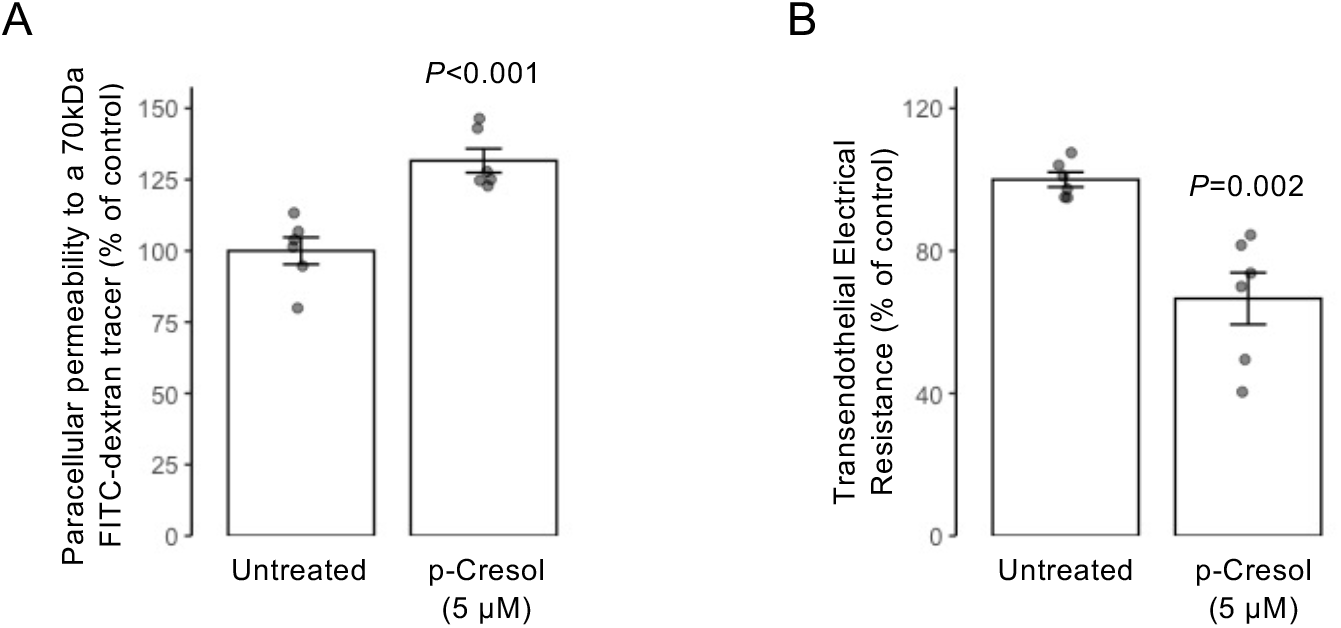
Treatment with *p*-cresol impairs endothelial barrier integrity *in vitro*. **A)** Paracellular permeability of polarised hCMEC/D3 monolayers to a 70 kDa FITC-dextran tracer following 24 h treatment with *p*-cresol (5 μM); data are mean ± s.e.m., n=6. **B)** Trans-endothelial electrical resistance across polarised hCMEC/D3 monolayers following 24 h treatment with *p*-cresol (5 μM); data are mean ± s.e.m., n=6.

**Supplemental Figure 2:**
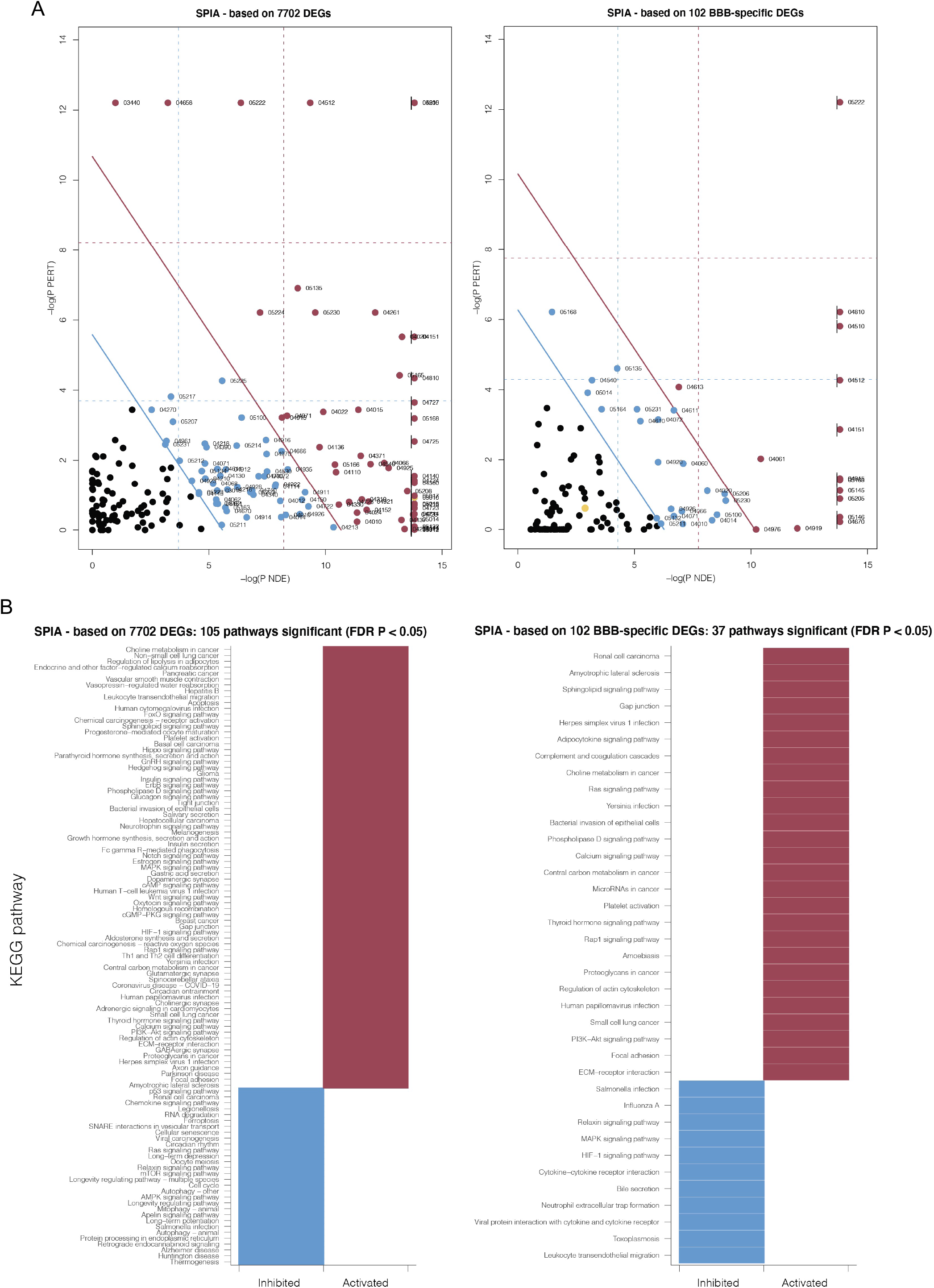
Signaling pathway impact analysis (SPIA) for gene expression in mouse brain cells following pCG treatment. **A)** SPIA results for all 7702 differentially expressed genes or for the 102 BBB-relevant differentially expressed genes in the CNS of male C57Bl/6 mice 2 h following i.p. injection of 1 mg/kg pCG (n=3 per group). The pathways in red to the right of the thick red line are significant after FWER correction of the global *P* values (pG, obtained by combining the pPERT and pNDE using Fisher’s method). The pathways in blue to the right of the thick blue line are significant after FDR correction of the pG values. Numerical labels refer to the KEGG pathway. **B)** Summary of above SPIA results, indicating which KEGG pathways were activated (red) or inhibited (blue).

**Supplemental Figure 3:**
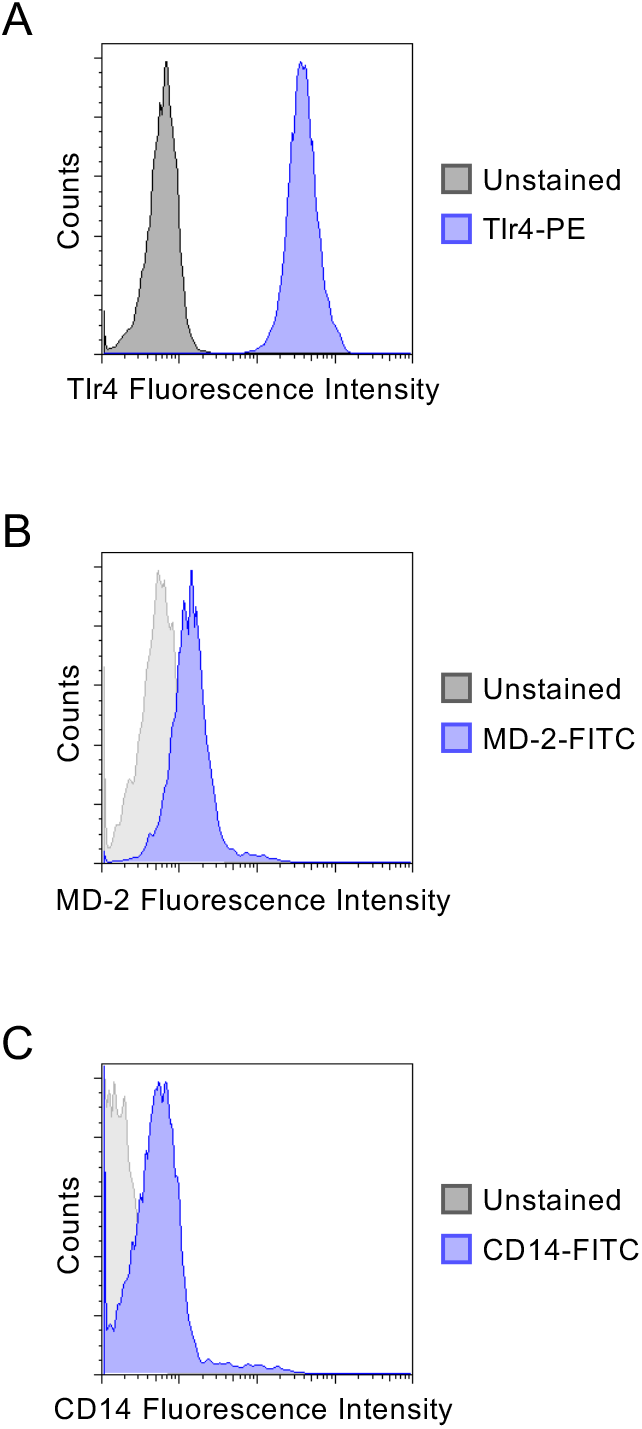
Expression of TLR4, MD-2 and CD14 by hCMEC/D3 cells. Typical flow cytometry histogram profiles of hCMEC/D3 cells immunolabelled with **A)** PE-conjugated anti-TLR4, **B)** FITC-conjugated anti-MD-2, or **C)** FITC-conjugated anti-CD14 antibodies.

**Supplemental Figure 4:**
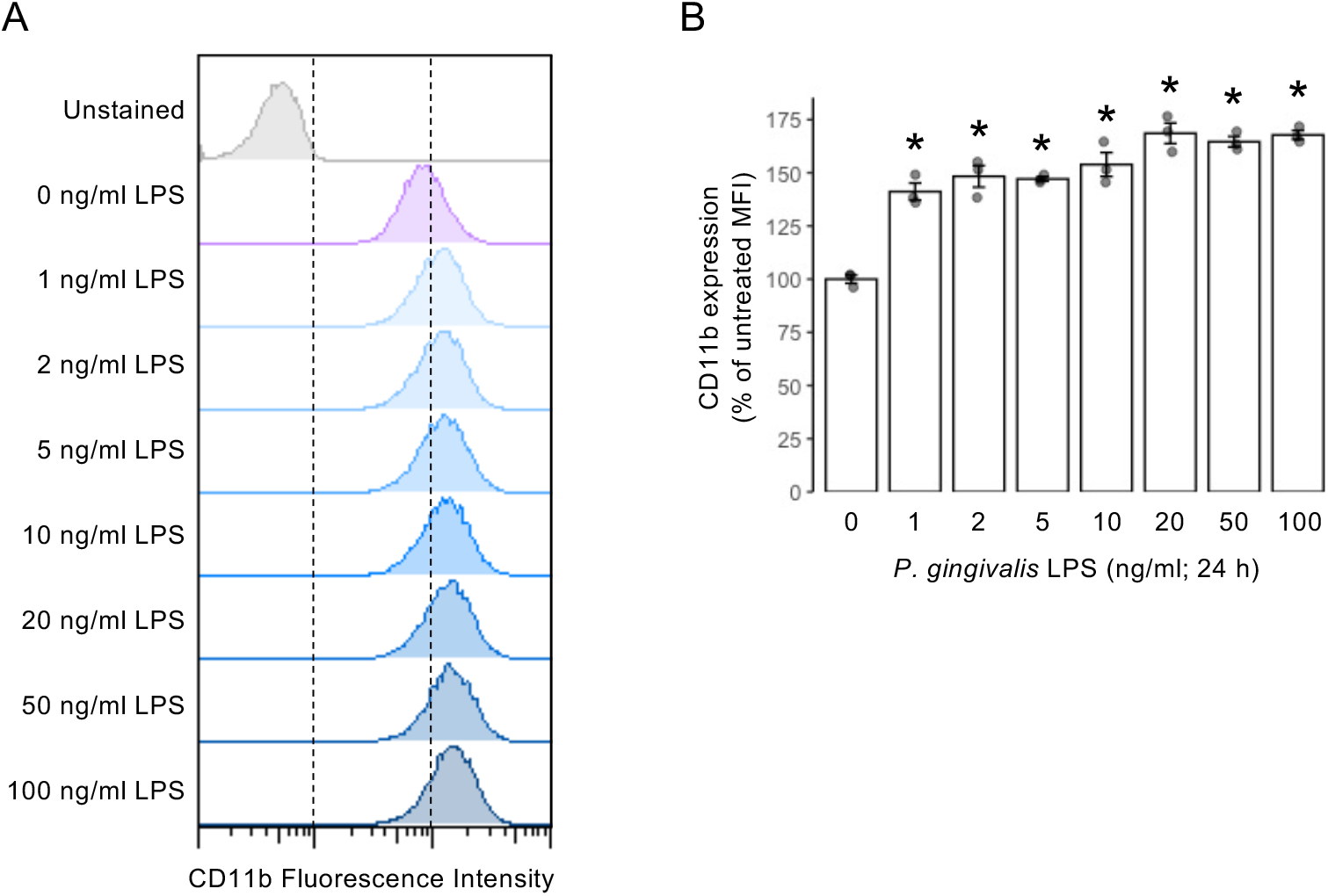
Exposure of THP-1 cells to LPS dose-dependently increases cell surface CD11b expression. **A)** Typical histograms showing a dose-dependent increase in CD11b fluorescence intensity. **B)** Median fluorescence intensities of THP-cell surface CD11b expression in THP-1 cells treated for 24 h with different doses of *Porphyromonas gingivalis* LPS; data are mean ± s.e.m., n=3, \**P*<0.05 vs. untreated cells.

